# Decoding Demosponge Diversity: Bayesian Analysis of Biodiversity, Extinction Events and Environmental Influences throughout the Phanerozoic

**DOI:** 10.1101/2025.01.24.634792

**Authors:** Astrid Schuster, Donald E. Canfield

## Abstract

Sponges (phylum Porifera) have been essential to marine ecosystems for over 600 million years, contributing to nutrient cycling, reef building, and ecological stability. Despite their evolutionary importance, the drivers of their diversity dynamics remain poorly understood. This study investigates the diversification patterns of Demospongiae, the largest class of sponges with a well-documented fossil record throughout the Phanerozoic Eon (541–0 Ma). Using fossil occurrence data and a Bayesian framework, we modeled origination and extinction rates to understand their evolutionary history.

Our findings reveal key extinction events, including three previously unrecognized events in the Cambrian, Late Silurian, and Late Jurassic, alongside known events such as the Permian-Triassic and Triassic-Jurassic extinctions. Notably, there was no statistical evidence for mass extinction during the Late Ordovician or Late Devonian. Additionally, Demosponges underwent a significant decline prior to the Cretaceous-Paleogene extinction.

To explore abiotic influences, we applied the Multivariate Birth-Death (MBD) model across all extinction events. This analysis identified correlations between major speciation and extinction events and key variables, including temperature, continental fragmentation, oxygen levels, and sea level, as well as geochemical proxies such as sulfur and strontium, which may reflect underlying drivers like anoxia, weathering, or tectonic activity. Temperature and oxygen levels, in particular, correlated with speciation and extinction during the Permian-Triassic extinction.

These findings underscore the role of environmental fluctuations in shaping demosponge diversity and highlight the complex interplay between changes in abiotic factors and sponge evolution. Our work provides critical insights into the factors driving sponge biodiversity and offers perspectives on how modern sponges might respond to ongoing climate change.

**Significance Statement:** Sponges are ancient and ecologically pivotal marine organisms that have persisted through Earth’s major environmental upheavals. Despite their resilience, the mechanisms driving their evolutionary responses to past climate shifts remain unclear. By leveraging the fossil record of Demospongiae, this study identifies previously unrecognized extinction events and elucidates how abiotic factors, including temperature and oxygen levels, shaped their diversity during these crises. These findings enhance our understanding of sponge evolutionary dynamics and resilience, offering critical insights into the vulnerability of marine ecosystems under current and future climate change scenarios.

## Introduction

Animal diversity has evolved over the last 540 millions years through the interplay between the emergence of new species and extinction. This diversity has been driven by both biotic factors, such as competition, predation and evolution, and abiotic factors, including climate change, volcanism, meteorite impacts and tectonic events (1–5). Major mass extinctions have reset the evolutionary stage, starting with a drop in diversity and followed by new clades often replacing earlier ones. These changes have allowed different animal groups to rise and dominate at different periods throughout Earth’s History (2, 6). For example, most of the early evolved so-called “Cambrian” fauna have been replaced by later-emerging more modern animal forms (7–10). Indeed, animal evolution has been punctuated by five major mass extinction events, and these have all impacted the subsequent history of animal evolution (11–13). Indeed, significant declines in biodiversity are increasingly viewed as precursors to systemic ecological destabilization. For example, biodiversity loss resulting from the Permian-Triassic extinction, led to severe ecological collapse due to the significant loss of biodiversity, which reduced functional redundancy in ecosystems (14).

One group that exemplifies both resilience and the complex patterns of evolutionary change is the sponges (phylum Porifera). They emerged approximately 800 million years (Ma) ago based on molecular clock evidence (15), with the first crown-group sponge fossil from the late-Ediacaran Period (around 551–539 Ma) (16). The discrepancy between molecular clock age estimates and the earliest fossil sponge record is thought to result from the absence of siliceous spicules in crown-group sponges (17). In the modern oceans, sponges play a critical ecological role by impacting marine food webs and nutrient cycling through their filtering activities (18–20). They also provided the first biological control on silica cycling in the oceans (21) and became the first builders of biological reefs, a role they have maintained throughout the Phanerozoic Eon (22). Despite many dramatic environmental changes since their evolution (see S1), sponges continue to thrive in today’s oceans, maintaining their ecological importance (23).

Of the four modern sponge classes –Calcarea, Hexactinellida, Homoscleromorpha, and Demospongiae– the latter is today’s most diversified sponge clade (23–25) and will be the focus of our palaeobiodiversity study. Notably, the Demospongiae class includes groups of sponges, such as certain tetractinellid and lithistid taxa (22, 26–29), that have a well-documented and continuous fossil record, making them an ideal group for studying biodiversity through time.

The general history of sponge diversity is well studied, and it is dynamic (see S1 for a literature summary). Studies have suggested that losses in shallow-water sponge diversity during mass extinctions were driven by cold intervals associated with major glaciation, as well as warming events related to increased greenhouse gas concentrations and the expansion of marine anoxia (30–32). Conversely, expansions and speciation events in sponges were driven by more stable, warm, and nutrient-rich conditions, particularly during periods of high sea levels and increased habitat availability in shallow marine environments (33, 34). Furthermore, of the “Big Five” previously recognized mass extinction events, the episodes of major diversity loss for demosponges, the Permian-Triassic mass extinction (PTME) and the Triassic-Jurassic mass extinction (TJME) were the most severe (35).

Despite these insights, and the potential for excellent fossil preservation in certain sponge groups (36), there has been no prior rigorous investigation into the palaeobiodiversity of sponges through geological time. Thus, while some paleontological studies (see S1 and (22, 27, 33, 37, 38)) have provided important insights into fossil sponge distribution and abundance, they primarily offer summaries limited to specific locations, specific sponge groups of interest, specific occurrences and specific time periods. Although earlier research documents the temporal distribution of sponges species and their occurrences in specific sites, it does not statistically examine rates of sponge origination or extinction, nor the relationships between sponge diversity and potential controlling abiotic factors. In addition, biases and gaps inherent in the fossil record have further complicated and hindered deeper analyses of these trends. While Fun et al. (39) provided a high-resolution summary of Cambrian to Early Triassic marine invertebrate biodiversity, this comprehensive work notably excluded sponges from its analysis, highlighting a gap in understanding how sponge biodiversity has fluctuated across geological time. In particular, the fundamental diversification dynamics, such as rate of origination and extinction, and how these dynamics might be coupled to potential driving factors like temperature, seawater chemistry, oxygen levels, and tectonic activity is urgently needed if we want to understand the history of sponge diversity. In addition, such insights could prove pivotal in predicting and mitigating the impacts of present and future environmental changes on sponge biodiversity and its impact on marine ecosystems.

To address this gap, we modeled the diversification dynamics of the class Demospongiae by examining extinction and speciation rates in relation to abiotic factors including temperature, seawater chemistry, oxygen levels, and tectonic activity. In our analysis, we used a Bayesian framework implemented in PyRate (40–42) to estimate diversity, extinction and origination rates from the fossil demosponge record and to investigate their correlation with abiotic factors. In detail, we first explore the overall Phanerozoic diversity of Demospongiae using fossil occurrence data from the Paleobiology Database (PBDB) and the literature. This analysis reveals the long-term pattern of sponge diversification history and highlights important stages of mass extinctions, including two that were previously unrecognized. We then estimate the origination, extinction and net (origination-extinction) rates to assess biodiversity losses. Finally, we use a multivariate birth-death model (PyRateMBD (43)) to evaluate the potential abiotic factors driving the extinctions and recovery of Demospongiae across both the known “Big Five” extinction events and the newly discovered extinctions.

## Results

### Diversification dynamics

Our Bayesian analysis reveals key diversification patterns of Demospongiae throughout the Phanerozoic, highlighting both well-known and newly identified extinction events that shaped their biodiversity (Fig. 1, S2).

**Figure 1.**
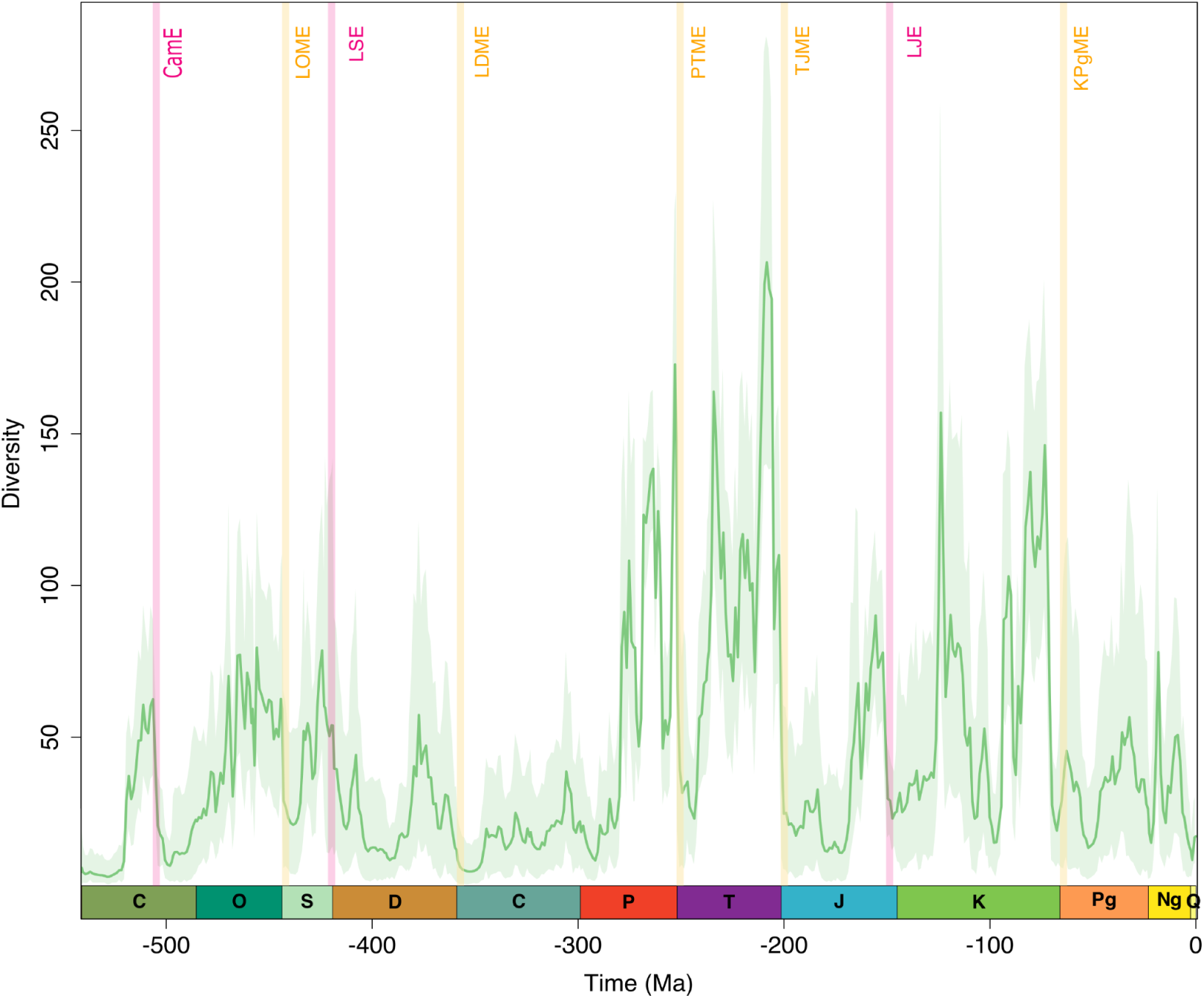
Large-scale trends of diversification dynamics of Demospongiae over the past ∼560 Ma. Displayed is the result of the mcmcDivE analysis from PyRate that allows to infer a corrected diversity trajectory across arbitrarily defined time bins (b=541 in this dataset). Dark, bold green line indicate the mean diversity with shaded areas indicating 95% highest posterior density (HPD) intervals. The ‘Big Five’ mass extinctions are indicated in light yellow: Late Ordovician mass extinction (LOME), Late Devonian mass extinction (LDME), Permian-Triassic mass extinction (PTME), Triassic-Jurassic mass extinction (TJME), and Cretaceous-Paleogene mass extinction (KPgME). Additional extinction events from this study are indicated in pink: Cambrian Extinction (CamE), Late Silurian extinction (LSE) event and the Late Jurassic extinction (LJE) event. C Cambrian, O Ordovician, S Silurian, D Devonian, C Carboniferous, P Permian, Tr Triassic, J Jurassic, K Cretaceous, Pg Paleogene, Ng Neogene, Q Quaternary.

Our results show a significant Cambrian extinction event (CamE, Fig. 1, Fig. 2a), which, while absent from the traditional “Big Five” mass extinctions in Earth history, is present in the demosponge record. This event marks a sharp decline in diversity, resulting in a negative net diversification rate (Fig. 2b) and a high extinction rate (Fig. 2a). Several genera, such as *Hazelia* (Cambrian: 61 occurrences, Post-Cambrian: 0 occurrences (occ.)), *Choia* (Cambrian: 52 occ., post-Cambrian: 1 occ.), *Leptomitus* (Cambrian: 43 occ., post-Cambrian: 0 occ.), *Paraleptomitella* (Cambiran: 34 occ., post-Cambrian 0) and *Vauxia* (Cambrian: 31 occ., post-Cambrian 0) which were once abundant during the Cambrian became extinct or nearly extinct (see genus *Choia*) by the end of the Cambrian Period (see S3).

**Figure 2.**
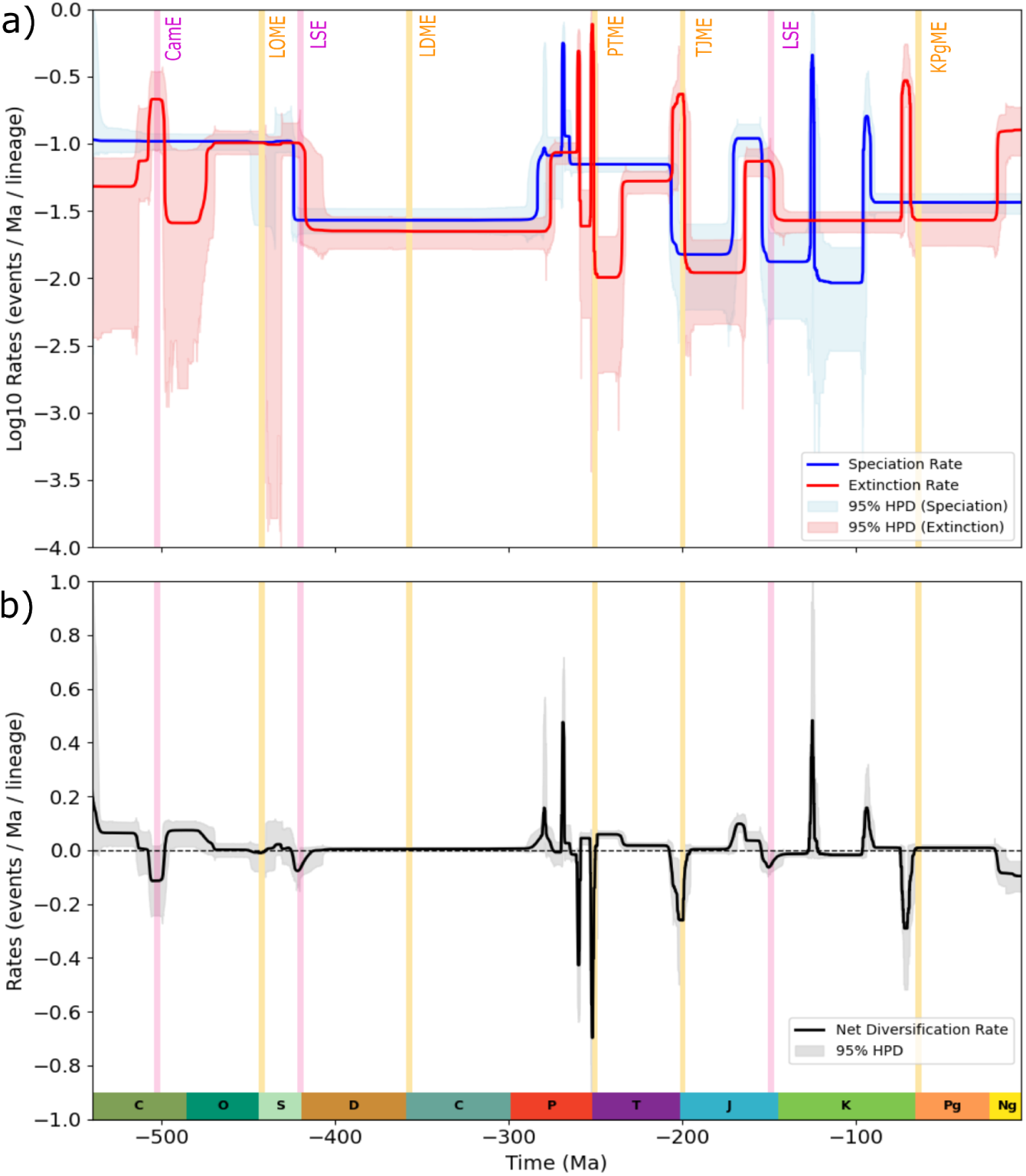
Rates of diversification of Demospongiae. a) Speciation and Extinction rate. b) Net diversification rate, calculated as speciation minus extinction rate. Shown are the mean values as solid lines and the 95% HPD intervals as shaded areas quantifying the uncertainty in rates. The ‘Big Five’ mass extinctions are indicated in light yellow and additional extinction events from this study are indicated in pink and labeling corresponds to Figure 1 of this study. C Cambrian, O Ordovician, S Silurian, D Devonian, C Carboniferous, P Permian, Tr Triassic, J Jurassic, K Cretaceous, Pg Paleogene, Ng Neogene, Q Quaternary.

Following the Cambrian Period, demosponges underwent a marked radiation during the early and mid-Ordovician, as evidenced by increasing diversity (Fig. 1). This pattern aligns with the Great Ordovician Biodiversity Event (GOBE), a period of significant taxonomic turnover that shifted marine ecosystems from the Cambrian to the Paleozoic faunal assemblages. Notably, our results indicate that this diversification occurred despite the constancy of speciation rates during the Ordovician (Fig. 2a). Instead, the observed Ordovician rise in diversity is primarily driven by a substantial reduction in extinction rates compared to the Cambrian extinction event (CamE), as evidenced by the sharp decline in the extinction curve during early and mid-Ordovician (Fig. 2a). This decoupling of speciation and extinction dynamics underscores the role of reduced extinction pressure in facilitating demosponge diversification during this interval.

Remarkably, our findings challenge the characterization of the Late Ordovician Mass Extinction (LOME) as a significant extinction event for demosponges, despite its status as the first of the “Big Five” extinction events. Although a decline in demosponge biodiversity is apparent during the Late Ordovician (Fig. 1), this pattern is not reflected in the underlying speciation and extinction rates (Fig. 2a), which remain consistently stable from the mid-Ordovician through the end of the Silurian. Biodiversity in the Silurian reached levels comparable to those observed in the mid-Ordovician (Fig. 1). However, this recovery was short-lived, as diversity began to decline towards the Silurian-Devonian boundary (Fig. 1, LSE). This Late Silurian Extinction event coincided with reductions in both speciation and extinction rates (Fgi. 2a), leading to fluctuations in net diversification (Fig. 2b), which ultimately turned negative by the end of the Silurian. Although our results indicate a LSE event for demosponges, it is not considered one of the “Big Five” mass extinction events.

Interestingly, no discernible signal for the Late Devonian Mass Extinction (LDME) is evident in our data as both speciation and extinction rates remained stable throughout this interval (Fig. 2a, 2b). Carboniferous diversity fluctuated but failed to recover to pre-LDME levels. The Permian-Triassic Mass Extinction (PTME) caused one of the most dramatic collapses in speciation and overall demosponge diversity (Fig. 1, 2a, b). This pattern is marked by two pronounced peaks in extinction rates (Fig. 2a), one occurring before the PTME and another during the PTME itself. Despite the substantial biodiversity loss associated with the PTME, overall diversity remained higher than the levels observed during the Devonian-Carboniferous transition (Fig. 1). Demosponge diversity peaked at its highest levels of the Phanerozoic during the late Triassic, with remarkably elevated diversity also recorded in the Permian (Fig. 1).

The Triassic-Jurassic Mass Extinction (TJME) caused a steep decline, evident in elevated extinction rates (Fig. 2b). Following this, our data indicate a Late Jurassic Extinction (LJE) event, marked by a decline in diversity and speciation rates (Fig. 1, 2a). Although less severe than the PTME and TJME before, the LJE contributed to persistently low diversity level that endured until the mid-Cretaceous period. The Cretaceous period experienced fluctuating diversity levels, leading to a significant decline shortly before the well-documented Cretaceous-Paleogene Mass Extinction (KPgME) (Fig. 1). Notably, extinction rates during this decline reached levels comparable to those observed during the TJME, preceding the KPgME (Fig. 1, 2a). Following the Cretaceous, net diversification stabilized at substantially lower levels, persisting throughout the Paleogene and Neogene periods (Fig. 1).

### Drivers of demosponge diversity

To evaluate the influence of abiotic factors and tectonic shifts on demosponge origination and extinction rates, we applied a Multivariate Birth-Death Model using *PyRateMBD* (43). Key extinction intervals analysed include the Late Ordovician Mass Extinction (LOME), Late Silurian Extinction (LSE), Late Devonian Mass Extinction (LDME), Permian–Triassic Mass Extinction (PTME), Triassic–Jurassic Mass Extinction (TJME), Late Jurassic Extinction (LJE) and Cretaceous–Paleogene Mass Extinction (KPgME) (Fig. 3). The Cambrian extinction (CamE) was excluded from this analysis due to insufficient predictor data available for this time period. For the extinction events analysed, we found no significant correlations for origination or extinction during the KPgME (Fig. 3a). During the LJE, strontium isotopes showed a negative correlation with origination (ω = 0.63), while continental fragmentation exhibited a positive correlation (ω = 0.65) (Fig. 3b). During the TJME, oxygen levels displayed a positive correlation with origination (ω = 0.87), and continental fragmentation showed a negative correlation with extinction (ω = 0.65) (Fig. 3c).

**Figure 3.**
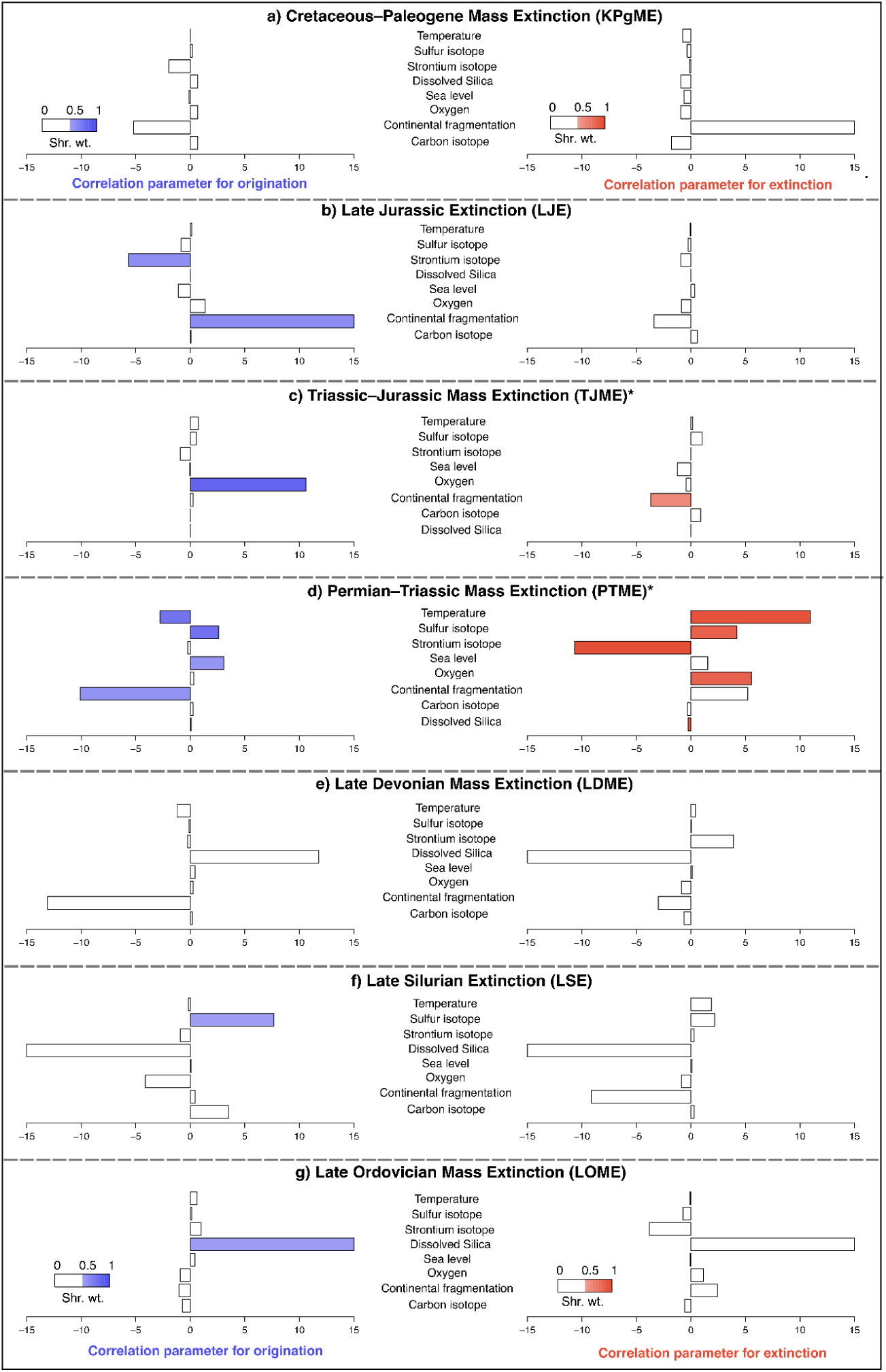
Estimated correlation parameters on origination (blue) and extinction (red) rates analyzed with the Multivariate Birth-Death model (PyRateMBD) with abiotic factors for the “Big Five” mass extinction (a, c, d, e, g) and two additional extinction events (b & f). A filled bar indicates a significant relationship (Shrinkage weight [Shr. Wt.] ω>0.5). Please note the different ordering of abiotic factor names in c) and d), marked with an asterisk (*), to match the corresponding PyRateMBD output .r and .log files.

The Permian–Triassic Mass Extinction showed a negative correlation between increasing temperature and origination (ω = 0.78) and a positive correlation with extinction (ω = 0.95) (S4). Sulfur isotopes correlated positively with both origination (ω = 0.80) and extinction (ω = 0.87). Strontium isotopes showed a negative correlation with extinction (ω = 0.97). Sea level fluctuations exhibited a positive correlation with origination (ω = 0.59) and no significant correlation with extinction (ω = 0.48). Oxygen levels correlated positively with extinction (ω = 0.87) but showed no correlation with origination (ω = 0.46). Continental fragmentation correlated negatively with origination (ω = 0.60). Dissolved silica displayed a positive correlation with origination (ω = 0.59) and a negative correlation with extinction (ω = 0.89) (Fig. 3d) (S4). No significant correlations for origination or extinction were observed during the LDME (Fig. 3e). During the LSE, sulfur isotopes correlated positively with origination (ω = 0.54) (Fig. 3f). During the LOME, dissolved silica showed a positive correlation with origination (ω = 0.55) (Fig. 3g).

## Discussion

Our findings reveal significant insights into the diversification dynamics of Demospongiae throughout the Phanerozoic Eon, particularly through the identification of previously unrecognized extinction events—the Cambrian Extinction (CamE), the Late Silurian Extinction (LSE) and the Late Jurassic Extinction (LJE) (Fig. 1, 2). Furthermore, our analysis reveals that certain major extinction events traditionally classified within the “Big Five”, including the LOME, LDME, and KPgME (Fig. 1), appear to have had no impact on demosponge diversity. Thus, this work provides a comprehensive framework for understanding the complex interplay between speciation and extinction rates across geological time and potential abiotic drivers.

Despite a decline in demosponge biodiversity during the Late Ordovician (Fig. 1), our analysis shows stable speciation and extinction rate from the mid-Ordovician through the end of the Silurian (Fig. 2a) suggesting that this constancy may result from a poorly known fossil sponge record as pointed out by the low sampling number in the dataset (see S5) rather than a pronounced extinction pulse. The lack of strong evidence for a major extinction event aligns with Sheehan (44), who noted that biodiversity declines during the Ordovician were likely driven by environmental changes rather than mass extinctions. Importantly, the impact of these changes varied depending on the environment, with biota in epicontinental seas strongly affected by sea-level declines, while organisms in open marine settings were less impacted. Since many demosponges inhabited deep-waters they may have shown more resilience to this extinction, unlike more shallow water sponges (stromatoporoids) which suffered greatly (45). Additionally, tropical biota experienced less-severe temperature changes compared to those at higher latitudes (44). Our PyRate MBD analysis (Fig. 3g) further identifies dissolved silica as a significant positive correlate with origination rates, consistent with Conley & Carey (46), who emphasized the importance of silica availability for marine biodiversity including sponges. During the Late Ordovician, a major glaciation event triggered the LOME, leading to a dramatic drop in sea levels and widespread environmental stress (e.g. 47). While this event disrupted marine ecosystems, our analysis suggests that demosponges were less impacted than other marine taxa, as indicated by the stable speciation rates observed. As the climate warmed following the glaciation, marine ecosystems began to recover, and the increased availability of silica may have played a crucial role in facilitating the diversification of marine taxa, including sponges, during this recovery phase.

The Late Silurian Extinction (LSE), identified in our study, corresponds with the well-documented Lau Event, a critical phase in demosponge evolution that has not been adequately addressed in previous research. This extinction is linked to significant environmental stressors, including high sea level rises and perturbations in the carbon and sulfur cycles, reflecting widespread anoxic conditions that likely disrupted marine ecosystems during the Late Silurian Period (48, 49). Most of the Silurian Period, however, is characterized by stable, warm climates, high sea levels, and only localized or minor episodes of anoxia (50), the fossil record suggests an increase in non-litistid demosponge diversity throughout the Silurian, primarily due to their diversification (51). However, our analysis reveals a marked decline in demosponge diversity following the LSE (Fig. 1). This decline is characterized by decreased speciation, elevated extinction rates, and a negative net diversification rate (Fig. 2a, b), which contrasts with the generally increasing sponge diversity reported in the literature (see S1). Notably, we observed a positive correlation between sulfur isotopes and origination rates during the LSE (Fig. 3f). An increase in δ^34^S is often related to increases in pyrite burial and the expansion of anoxia (52, 53). Indeed, expanded anoxia during the LSE has been previously correlated with extinction at this time (54). A positive relationship between δ^34^S and diversification is more difficult to explain, but could be related to elevated primary production associated with phosphorus release during anoxia (55), or mismatches in the temporal resolution of the sulfur isotope and diversity records (see S6). Interestingly, unlike tabulate and rugose corals, which experienced biodiversity declines likely driven by sea-level fluctuations and anoxic events (e.g. (56, 57), our analysis did not identify any extinction-related parameters correlated with these stressors in demosponges. This suggests that sponges may have exhibited greater resilience to such environmental perturbations during the Late Silurian. However, this resilience should be interpreted cautiously, due to the limited temporal accuracy of the sulfur isotope record (58) (S6), which may not adequately capture short time periods like the LSE.

The Devonian Period ended with a mass extinction, yet our analysis does not reveal a discernible signal of this event (Fig. 2). The Late Devonian anoxia (59) and glaciation, which contributed to the Late Devonian Mass Extinction (LDME) (60), are not evident in our PyRateMBD analysis of environmental predictors, nor are any other factors (Fig. 3). This suggests that sponges, despite the challenging environmental conditions of the time, may have been relatively unaffected by these perturbations compared to other marine taxa such as brachiopods (61), rugose and tabulate corals (62), and calcite-based stromatoporoid sponges (63) that suffered greatly.

The observed reduction in sponge biodiversity (Fig. 1) does not coincide with the LDME but instead appears to precede it. This suggests that environmental conditions may have already deteriorated for sponges prior to the extinction event itself, potentially driving a decline in diversity before this event. By the onset of the Late Devonian extinction, sponge diversity was already at an all-time low, which may explain why sponges were less impacted by the extinction drivers that affected other animal groups. This pattern may reflect a broader ecological sensitivity of sponges to prolonged environmental stress, at least to certain types of decline, rather than to acute extinction events. A similar trend is observed at the end-Cretaceous extinction (see discussion below), which further supports the notion that sponges might respond differently to long-term environmental changes compared to other taxa. These dynamics underscore the need for further investigation into how sponges are affected by both gradual environmental deterioration and abrupt extinction drivers.

Our analysis also highlights the evolutionary trajectory of demosponges during the transition from the Devonian to the Carboniferous Period, with taxa such as *Hindia* and *Pachythecata* giving way to forms resembling modern Mesozoic groups, including *Chaetetes*, *Haplistion*, and the Chaetetidae (S7). The higher prevalence of lithistid sponges in the Devonian compared to the Carboniferous (S7) suggests a transition from lithistid-dominated to non-lithistid sponge communities, aligning with findings from Vishnevskaya et al. (64). Despite this observed transition, our findings indicate that the LDME, one of the “Big Five” mass extinctions, did not have a profound impact on the origination or extinction of demosponges.

In contrast, trends in sponge biodiversity through the PTME show major mass extinctions, including a two stage extinction (Fig. 2a), thus reflecting broader patterns of mass extinction as shown through biodiversity patterns of a broad range of animal groups (6, 14, 56, e.g. 61, 65–68). Our Multivariate Birth-Death Model (PyRateMBD) (Fig. 3) reveals that the drivers of demosponge diversity during the Permian-Jurassic period strongly correlate with abiotic factors influencing both speciation and extinction rates. Specifically, increasing temperatures are significantly associated with both a negative origination and a positive extinction correlation (Fig. 3d), suggesting that thermal regimes played a crucial role in shaping demosponge diversification during this critical interval by reducing speciation and causing a higher extinction rate (Fig. 3d). This observation aligns with Song et al. (69), who identified temperature excursions of ∼3°C above the long-term average that trigger mass extinctions, emphasizing that exceeding a 5°C increase significantly elevates extinction probabilities. Similarly, coral studies show that bleaching and mortality rates exceed 60% when sea surface temperatures rise by these amounts (70).

These patterns indicate a sensitivity of marine communities, including Demospongiae, to temperature fluctuations. Moreover, the 485-million-year history of Earth’s surface temperature presented by Judd et al. (71) underscores the long-term influence of thermal dynamics on marine biodiversity, highlighting that significant temperature fluctuations, particularly during major mass extinction events such as the PTME and TJME, have historically driven extinction and speciation events, with documented losses of up to 90% in some marine environments (72). Moreover, studies indicate that shifts in sea temperature and acidification have historically driven changes in coral diversity (56, 73, 74), much like the thermal regimes impacting demosponge communities during the Permian–Jurassic period. This sensitivity of animal diversity to rapid increasing temperatures aligns with our findings that demonstrate significant correlations between increasing temperatures and both negative origination and positive extinction rates in Demospongiae (Fig. 3d).

Sulfur, released by massive volcanic eruptions and the breakdown of organic matter in anoxic conditions, contributed to the formation of hydrogen sulfide (H₂S), a toxic compound that can be lethal to aerobic organisms (Erickson et al., 2020). This sulfur release, particularly in shallow marine environments where many sponge species reside, has been implicated in the PTME (Kump et al., 2005). The accumulation of H₂S in these regions likely disrupted marine ecosystems by reducing oxygen availability and through sulfide toxicity, leading to a stressful environment for aerobic taxa. Our analysis also identifies a positive correlation between sulfur isotopes and extinction rates (Fig. 3d). Elevated δ^34^S can be associated with increased pyrite sulfur burial under anoxic conditions. Therefore, our data are consistent with significant extinction pressures imposed by sulfur-driven anoxia (75). We also observe a positive correlation between sulfur isotopes and origination rates (Fig. 3d). This correlation is harder to explain, but as discussed above for the LSE, this correlation could be related to increased productivity associated with phosphorus release under anoxic conditions, or a mismatch in the temporal resolution between the sulfur isotope and extinction records.

The overall recovery of sponge biodiversity, however, remained slow following major extinction events such as the LSE, TJME and LJE, reflecting the lingering impacts of anoxic conditions. In contrast, after the PTME, sponges exhibited a relatively rapid recovery compared to other taxa (76), likely due to their resilience to low-oxygen environments and ability to thrive under conditions unfavorable to many other marine organisms. Strontium isotopes are sensitive weathering indicators, where increasing values are correlated with increasing weathering intensity, and decreasing values with reduced weathering intensity, where weathering intensity controls nutrient input to the oceans (77, 78). Thus, a negative correlation between strontium isotopic composition and extinction rate (Fig. 3d) would imply that reduced nutrient availability could have been an additional stressor adding to the severity sponge biodiversity loss during the PTME.

The positive correlation between origination rates and sea-level rise following the PTME suggests that the subsequent expansion of shallow marine habitats may have facilitated sponge speciation (Fig. 3d). Rising sea levels likely increased the availability of suitable ecological niches (79), promoting diversification by offering new areas for sponge colonization. The fragmentation of the Pangea supercontinent created new oceanic and terrestrial barriers, disrupting previously continuous marine ecosystems and isolating populations (5). For sponges, this geographic isolation likely limited gene flow between populations, reducing genetic diversity and the potential for speciation as evident by our negative speciation parameter (Fig. 3d). However, despite the strong correlations observed, the actual changes in the fragmentation index are relatively small, suggesting that continental fragmentation may have had a more limited direct impact on sponge origination during this period. Nonetheless, the formation of physical and ecological barriers constrained the expansion of sponge habitats, thereby limiting the availability of new ecological niches required for adaptive radiation. These conditions were particularly unfavorable for the origination of new demosponge species, as the combined effects of restricted dispersal, habitat fragmentation, and heightened environmental instability posed substantial challenges to diversification (80), as evident from our analysis (Fig. 2, 3d).

Silica (SiO₂) availability during the PTME appears to have been relatively stable or even favorable, as evidenced by the slightly negative correlation with extinction and the slightly positive correlation with origination (Fig. 3d). Sponges utilize silica to construct their spicules, a critical structural component that provides support and defense (21, 81, 82). This stability in silica availability likely supported the persistence of silica-dependent demosponges such as lithistids, enabling certain taxa to endure or recover more effectively in the aftermath of the PTME. The modest positive correlation with origination (Fig. 3d) suggests that some new sponge species capable of utilizing silica emerged during this period, though their diversification may have been constrained compared to more adaptable or resilient groups.

In contrast to the PTME, oxygen shows a positive correlation parameter for origination at the TJME (Fig. 3c) which suggests that localized, oxygenated refugia may have supported new species despite widespread oceanic hypoxia (83). Shifts in ocean currents or stratification likely created temporary areas of higher oxygen, enabling sponge species to thrive and diversify in these niches. The negative correlation between continental fragmentation index and extinction during the TJME (Fig. 3c) suggests that the breakup of Pangea may have created refugia or isolated environments where sponge populations could persist. This fragmentation could have reduced the impact of global environmental stressors by creating more stable, localized habitats, which may have buffered sponge species from the broader extinction pressures (73, 84).

The Late Jurassic Extinction (LJE) represents a significant but underappreciated decline in demosponge diversity (Fig. 1) during a period typically considered climatically stable (5, 85). Characterized by reduced speciation and elevated extinction rates, resulting in a negative net diversification rate (Fig. 2), this event coincided with the breakup of Pangea. Although not classified as one of the “Big Five” mass extinction events, evidence of biodiversity loss in other marine taxa, such as corals, underscores its ecological importance (73). Indeed, coral reef decline across the Jurassic-Cretaceous boundary are linked to anoxic conditions, sea-level fluctuations, and shifts in ocean chemistry (86–88). Our analysis identifies a negative trend in strontium isotopes for origination (Fig. 3b), which likely reflects reduced nutrient input (including Si) to the oceans that hindered spicule formation, a critical structural component for sponges (82). Conversely, the positive correlation with continental fragmentation suggests that the breakup of Pangea created new ecological niches, fostering diversification in isolated habitats (80, 84). This interplay of factors illustrates the complexity of sponge biodiversity dynamics during this transitional period.

Our biodiversity analysis (Fig. 1) reveals a significant decline in demosponge diversity preceding the official timing of the KPgME. This early drop is further corroborated by elevated extinction rates (Fig. 2a) and a negative net diversification rate (Fig. 2b). This pattern raises the question of why demosponge diversity declined well before the KPgME, an event characterizing critical impacts on other animal groups such as brachiopods and bivalves (e.g. 61). We could not identify any clear correlation factors for this pre-event decline (Fig. 3a), which is however, not surprising given that the sponge extinction predates the broader KPgME disruptions. This temporal disconnection aligns with carbon isotope measurements across the K-T boundary which reveal severe, rapid and repeated fluctuations in oceanic productivity during the 3 m.y leading up to the final KPgME event (89, 90). Additionally, productivity and ocean circulation remained suppressed for at least several tens of thousands of years following the boundary (91), further highlighting the prolonged environmental instability of this period. Therefore, these changes could have devastated marine ecosystems before the KPgE potentially contributing to pre-event declines in species richness. Nevertheless, this anomaly warrants further additional biotic exploration towards species competition or predation, which could offer insights into this enigmatic extinction event in sponges.

In conclusion, this study offers a comprehensive exploration of the diversification dynamics of Demospongiae throughout the Phanerozoic, identifying key extinction events that were previously underrecognized, including the Cambrian Extinction (CamE), Late Silurian Extinction (LSE), and Late Jurassic Extinction (LJE) (Fig.1 & S2). Our analysis highlights that certain major extinction events, notably those within the “Big Five,” did not impact demosponge biodiversity, challenging previous assumptions about the role of these mass extinctions in shaping sponge diversity. We further identified significant shifts in extinction and speciation rates (Fig. 2), offering a comprehensive framework for interpreting past biodiversity changes. The findings emphasize the sensitivity of sponge communities to global climate events and provide crucial insights for predicting future biodiversity trends amidst accelerating climate change. These results not only contribute to understanding the historical resilience of sponge ecosystems but also offer a predictive tool for monitoring biodiversity responses to ongoing environmental shifts.

## Materials and Methods

### Fossil occurrence data

The long-term pattern of diversification dynamics of the class Demospongiae was the focus of this study, as this class is well represented in the fossil record throughout the Phanerozoic Eon (541–0 Ma). Fossil occurrence data were downloaded from the Paleobiology Database (PBDB) on 09/01/2024 (http://paleobiodb.org/). Entries with uncertain genus or species names (cf., aff., ?, informal, ex.) were eliminated and only extinct taxa and accepted names were included. However, open species names (i.e. if the species name was sp.) were retained in the dataset. Further curation of the data included the check of species names for typos and inconsistent spelling by an algorithm integrated in PyRate (v.3) (40). Given our interest in the overall biodiversity of the class Demospongiae, all analyses were conducted at the class level. Thus, 21 occurrences of the class Demospongiae, 41 of the subclass "Lithistida," and 2 of Ceractinomorpha, along with 3 occurrences of the order Tabulospongida, 1 of Axinellida, 1 of Hadromerida, 1 of Spirophorida, and 9 of Chaetetida, were retained in the dataset following manual verification of their status and rank. In addition, a total of 117 occurrences were assigned an accepted family rank. Notably, out of the total 8325 occurrences, the majority were identified at the accepted species level (5972 occurrences) or genus level (2139 occurrences). In total, our dataset included 1,641 taxa ranging from 599.9 (+/-22.78) to 0.001 (+/- 0.001) Ma of which 743 have a single occurrence.

### Estimation of diversification rate

Diversity patterns are a result of origination and extinction processes, meaning a decrease in diversity may have resulted from elevated extinction rate or decreased speciation rate or both, and correspond to different mechanisms. Therefore, separating extinction and origination processes is necessary to calculate past biodiversity and investigate drivers of diversification (42). In this study, diversification dynamics were analyzed from the above dataset of Demospongiae using a Bayesian approach as implemented in PyRate (v.3) (40–42). Unlike other methods that calculate rate changes at separate, distinct time points (normally stage boundaries), PyRate estimates the preservation rates (*q*), the times of origination and extinction (*Ts* and *Te*) of each taxa and the origination and extinction rates (ለ and *µ*) by incorporating both the fossil preservation process and the birth-death process, while perceiving the diversification of a given group as a constant process. The fossil preservation process accounts for the unequal likelihood that species will be preserved in the fossil record, as not all species or environments favor fossilization. The birth-death process models how species originate (ለ) and go extinct (*µ)* over time, providing a continuous estimation of diversification patterns. Using the reversible-jump Markov Chain Monte Carlo (rjMCMC) algorithm (-A 4 option in PyRate), we estimated the number, timing, and statistical significance of changes in origination and extinction rates. This algorithm enables more accurate and reliable detection of biodiversity dynamics by explicitly accounting for the variability in fossil preservation and species turnover over geological time (42).

Additionally, we performed a maximum likelihood modeltest (-PPmodeltest option) as described in (42) to find the best-fit preservation process among the homogeneous Poisson process (-mHPP option, the non-homogeneous Poisson process (default option), and the time-variable Poisson process (TPP, -q option). The result of the model test supported the TPP process as the best fitting model for our data, which permits the rate of preservation to fluctuate in a series of time bins, each with a constant rate. For our dataset, we also accounted for varying preservation rates across taxa using the Gamma model (-mG option), with gamma-distributed rate heterogeneity. This method models the natural variability in preservation rates among different species by assuming that these rates follow a gamma distribution, which captures the uneven likelihood of fossilization across taxa. The shape parameter was set to 1.5 and the rate parameter was set to 0, allowing PyRate to estimate the rate directly from the data (pP 1.5 0). This approach enables us to account for taxon-specific differences in preservation, ensuring a more accurate estimation of our diversification dynamics.

We ran PyRate for 250,000,000 MCMC generations with a sampling frequency of 25,000, discarding the first 20% as burn-in. Effective sample size (ESS) was assessed using Tracer (v.1.7.1) (92) and was set to >200 for all parameters. Mean times and 95% highest posterior density intervals of origination, extinction and net diversification (origination minus extinction) were calculated from the combined log files (-combLogRJ option) from the 10 replicates and plotted using the -plotRJ function in Pyrate, set to -root_plot 541 -min_age_plot 0 -grid_plot 0.1 on a log scale (logT 1).

### Diversity estimation

For the diversity estimation we used two methods: the Bayesian model-based mcmcDivE (93) and the lineage through time (-ltt) function of PyRate. The first method takes the uneven sampling of the fossil sponge record into account and uses the preservation rates generated by PyRate and the occurrence data to estimate sampling-corrected diversity (93). Our mcmcDivE analysis was run for 50,000,000 iterations, sampled at every 50,000th iteration on the combined pyrate (-combLog) preservation log files. Convergence (EES >200) was assessed using Tracer (v.1.7.1) (92). The first 10% of the posterior samples were discarded as burn-in. The median and 95% highest posterior density intervals of diversity were calculated (Figure 1). The mean times for the lineage through time plot (Supplementary Figure 1) were generated using the -ginput function in PyRate after a 10% burn-in, and then plotted using the -ltt function with -grid_plot 0.1.

Relative abundance plots (S3 and S7) were generated using RStudio and were based on the lineage diversity output file from PyRate. All underlying data for this manuscript including all R and python scripts as well as PyRate generated files to run the R Shiny app are available in Zenodo (https://doi.org/10.5281/zenodo.14736214).

### Selection of potential drivers of diversification dynamics

In order to investigate potential abiotic drivers that might have influenced sponge biodiversity around the “Big Five” extinction events as well as during the discovered LSE and LJE, we selected eight predictors. Dissolved Silica (SiO_2_) (94), marine carbon isotope composition (δ^13^C) (95), global continental fragmentation (5), oxygen (96, 97), sea level (85, 98, 99), strontium isotope composition (^87^Sr/^86^Sr) (100), sulfur isotope composition (δ^34^S) (58) and sea water temperature (101) with their global average temperature estimated from δ^18^O. These factors have been used to test and explain diversification dynamics of other marine organisms such as brachiopods, bivalves (61) and bryozoans (102), but have never been tested for sponges.

### Multivariate birth-death analyses

The multivariate birth-death (MBD) model available in the software package PyRateMBD (43) was used to assess the influence of abiotic factors on the diversification dynamics of the class Demospongiae. The MBD model correlates time continuous variables, like in our case the abiotic factors of SiO_2_, δ^13^C, global continental fragmentation, oxygen, sea level, ^87^Sr/^86^Sr, δ^34^S, sea water temperature, with origination and extinction rates through an exponential or linear function. It uses the MCMC algorithm and estimas the baseline origination (ለ*_0_*) and extinction (µ*_0_*) rates, and correlation parameters (*G*ለ *and G*µ). A positive *G* implies that the factor correlates positively with the rate and vice versa. To prevent over-parameterisation, the analysis uses a horseshoe prior, which is a type of shrinkage prior specifically designed to handle situations where a large number of predictors are considered, but only a few of them are expected to have a significant effect. The horseshoe prior reduces the impact of irrelevant predictors by shrinking their correlation parameters towards zero, helping to prevent overfitting and maintain robust parameter estimates. A correlation parameter with a shrinkage weight (ω) close to 0 is likely to be noise, whereas a value close to 1 represents a true signal (43). In this study, a parameter weight threshold of 0.5 was used to distinguish significant correlations.

We performed this analysis for the class Demospongiae and for each extinction event separately. All analysis ran for 35,000,000 generations to ensure comprehensive sampling, from which 35,000 posterior samples were saved, representing the converged results for reliable statistical inference. To avoid re-modelling of the complex preservation analysis, the Tss and Tes estimates from the previous PyRate analyses were used as input with 10 replications. Convergence was assessed using Tracer (v.1.7.1) (92) and an ESS >200. The highest posterior density intervals (HPD 95%) of the correlation parameters and the shrinkage weights [Shr. Wt.] were calculated. A shrinkage weight of >0.5 meaning the relationship was significant (43). We analyzed both exponential (-m 0) and linear (-m 1) models for our dataset. In the exponential model, the rates of speciation and extinction are represented as exponential functions of a variable that changes continuously over time. On the other hand, the linear model presumes a linear relationship. The two models were assessed by calculating the Bayes factor following Lehtonen et al. 2017. In our case the linear model (-m1) was supported therefore only the result of the linear model is shown.

## Acknowledgments

We want to thank all the contributors of the PBDB. AS wants to thank D. Silvestro for assistance with PyRate. This project was funded by Villum Foundation grants no. 16518 and 54433.

## Supplementary Information S1

### Literature review of the fossil demosponge diversity and environmental influences.

Fossil sponges show some general biodiversity patterns among the class Demospongiae, revealing possible relationships between sponge diversity, climate, and ocean chemistry. For example, during the Cambrian Period, climates transitioned from cool (103, 104) to potential greenhouse conditions (105), with localized anoxia negatively affecting marine biodiversity including sponges (106). Demosponge clades like tetractinellids, halichondrids, and ceractinomorphs emerged during this time and were highly developed and taxonomically dependent on substrates and on tropical conditions (28, 107). By the mid-Cambrian Period, rigidly skeletonized demosponges known as lithistids dominated the fossil record, contributing to the early establishment of key demosponge faunas (22, 26, 108). Conversely, the once-abundant archaeocyathid reefs declined significantly by the late Cambrian Period (109, 110).

The Ordovician Period started warm but cooled, leading to major glaciation in Gondwana and to the Late Ordovician Mass Extinction (LOME) (111). This extinction, considered the second most severe of the entire Phanerozoic Eon (35), may have resulted from a combination of cooling and ocean anoxia (50). Lithistid demosponges thrived in shallow waters, especially during the Great Ordovician Biodiversification Event (GOBE) (112), although they were heavily impacted by the LOME (30). Deep-water siliceous sponges, including demosponges, showed resilience to this extinction, unlike the reef-forming stromatoporoids which suffered greatly (45).

The Silurian Period had stable, warm climates with high sea level and minor anoxia (50). Silurian reefs were dominated by stromatoporoids (113) and lithistid sponges (114) contributing to patchy reef formations (115). The fossil record indicates a steady increase in non-lithistid demosponge diversity through the Silurian Period, despite minor extinction events primarily affecting planktonic graptolites (116).

In the Devonian Period, initial warmth gave way to cooling and then to later warming, possibly influenced by plant evolution (117). The Devonian Period ended with significant anoxic events (59) and a glaciation that contributed to the Late Devonian mass extinction event (LDME) (60). During this early time period of the Phanerozoic, demosponges had evolved from ancient Paleozoic taxa to forms resembling modern groups. Stromatoporoids formed significant reefs with corals until they nearly vanished in the LDME (63). Meanwhile, deep-water siliceous sponges transited from lithistid-dominated to non-lithistid forms (64).

The early Carboniferous Period was warm but saw increasing glaciation and aridity towards its end, with periodic deep-water anoxia (118). This Period is known for diverse siliceous sponges, including the first freshwater demosponges reported in the upper Carboniferous (119). These sponges, along with sphinctozoans, occupied various habitats, though they were less common than during the Devonian Period (120). The Permian Period warmed from previous icehouse conditions, ending in extreme warming and widespread anoxia linked to volcanic activity, leading to the Permian-Triassic Mass Extinction (PTME), the most severe in the Phanerozoic (67, 121). Hypercalcified sponge groups dominated, but siliceous demosponges were also quite diverse at the beginning of the Permian (29). The PTME, however, led to the extinction of many sponge groups and a significant shift in demosponge biodiversity from Paleozoic to Mesozoic faunas (28, 122).

The Triassic Period began with severe greenhouse conditions and fluctuated through humid episodes, with anoxia delaying recovery from the PTME and contributing to the Triassic-Jurassic Mass Extinction (TJME) (83, 123, 124). The TJME led to the extinction of several sponge groups, but those that survived helped build a diverse reef ecosystem in the Jurassic Period (125). The warm conditions continued into the Jurassic, which was marked by dry low latitudes, expanding shallow seas, and persistent anoxia affecting marine life. In the early Jurassic demosponges and hexactinellids were abundant, with lithistids and stromatoporoids becoming key reef builders (125). A decline in siliceous reef-building sponges including lithistids, occurred later in the Mesozoic Era, likely due to oceanic silicon limitations (81).

The Cretaceous Period featured an extreme greenhouse climate, high sea levels (126, 127), and multiple Oceanic Anoxic Events (OAEs) (128, 129) driven by volcanism and oceanic changes, followed by cooling and significant sea-level shifts. During the Cretaceous Period, European regions were hotspots for diverse demosponge populations. Lithistids, particularly rhizomorines and dicranocladines, thrived in shallow seaways but declined towards the end of the Cretaceous (22, 130). Stromatoporoids became extinct at the Cretaceous–Paleogene Mass extinction (KPgME) (131), though the impact on other sponges is less well documented.

The Cenozoic Era started with the Paleocene-Eocene thermal maximum, associated with transient anoxia, then gradually cooled, leading into the current Ice Age with Arctic and Antarctic glaciation (132–134). This Era reveals a rich diversity of demosponges, particularly lithistids, with complex Eocene faunas (135–138), that is still ongoing research. Sponge diversity during the Cenozoic is thought to be more influenced by sea level changes than climate, with large biotas emerging during periods of high sea levels (33).

This short literature review highlights the complex interactions between climate, ocean chemistry, and biodiversity and illustrates how demosponges adapted and evolved through major extinction events and environmental fluctuations.

**Supplementary Figure S2:**
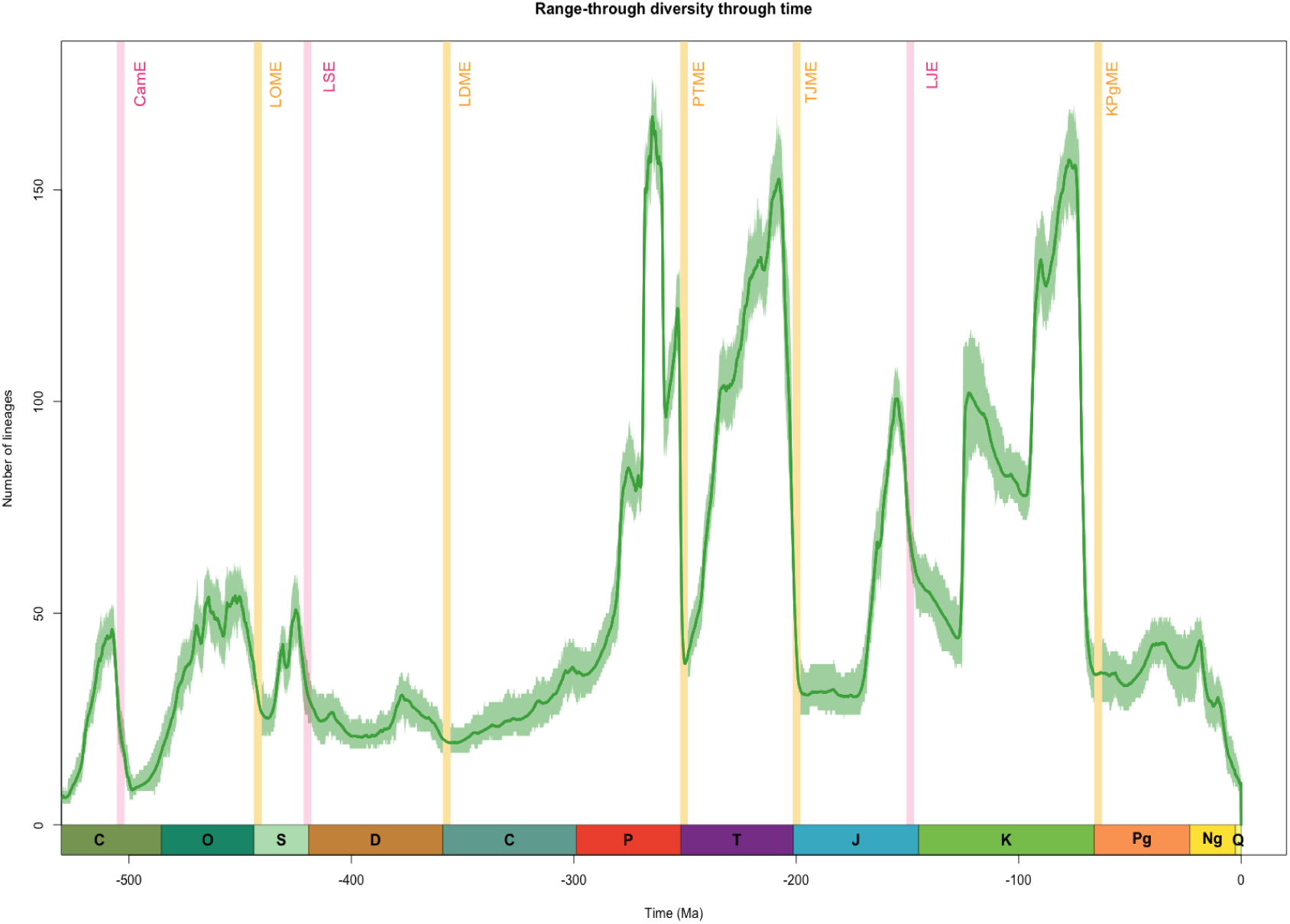
Large-scale trends of diversification dynamics of Demospongiae over the past ∼560 Ma. Displayed is the result of the lineage through time plot (-ltt function) in PyRate. Dark green line indicates the mean diversity with shaded areas indicating estimations of different replications that incorporate age uncertainties of the fossil occurrences. The ‘Big Five’ mass extinctions are indicated in light yellow: Late Ordovician mass extinction (LOME), Late Devonian mass extinction (LDME), Permian-Triassic mass extinction (PTME), Triassic-Jurassic mass extinction (TJME), and Cretaceous-Paleogene mass extinction (KPgME). Additional extinction events from this study are indicated in pink: Cambrian extinction (CamE), Late Silurian extinction (LSE) event and the Late Jurassic extinction (LJE) event. C Cambrian, O Ordovician, S Silurian, D Devonian, C Carboniferous, P Permian, Tr Triassic, J Jurassic, K Cretaceous, Pg Paleogene, Ng Neogene, Q Quaternary.

**Supplementary Fig. S3.**
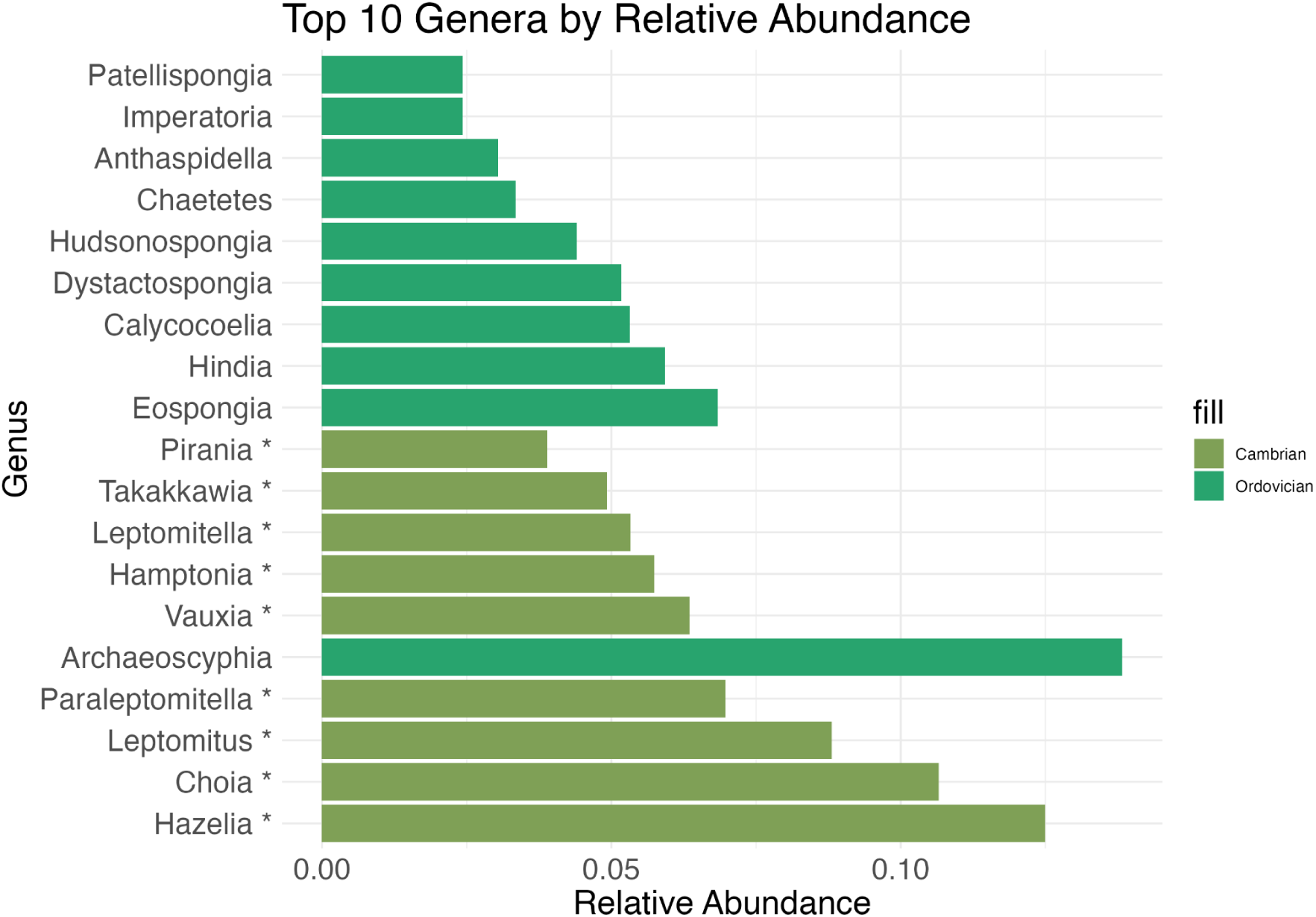
Relative abundance of top 10 dominant genera in the Cambrian and Ordovician Period. The plot shows the proportion of each genus within these two periods. Genera that became extinct in the Cambrian are marked with an asterisk. The relative abundance of each genus was calculated by dividing the count of occurrences by the total number of occurrences within the respective period, providing a normalized measure of each genus’s prevalence.

**Supplementary Table S4.**
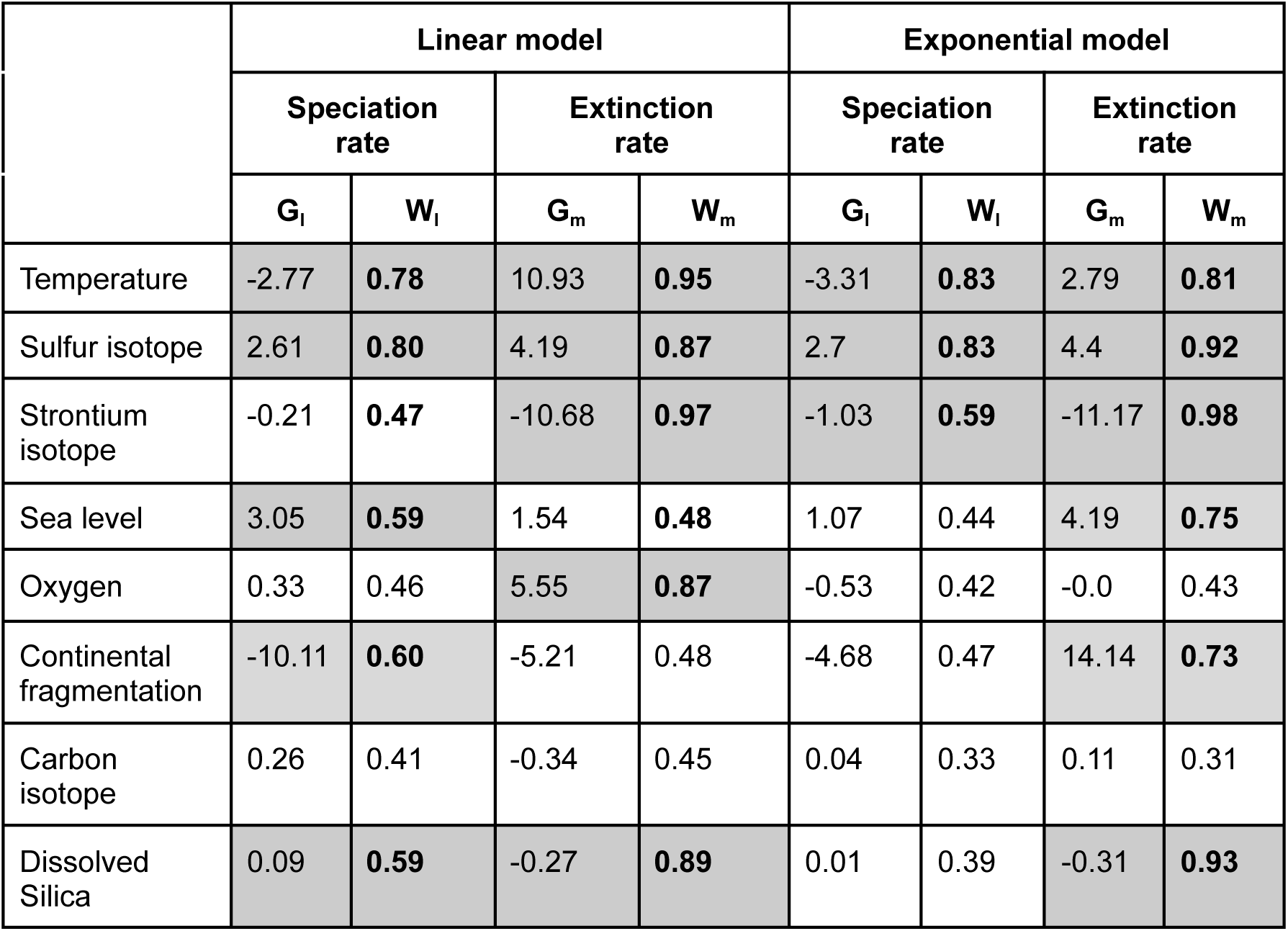
Correlation parameters (G_I_) and Shrinkage weight [Shr. Wt.] (W_I_) of the linear and exponential models with the occurrence dataset. Shown are results for the Perminan–Triassic, 272–227 Ma. A Shr. Wt. greater than 0.5 (WI>0.5) is highlighted in bold and indicates significant evidence for correlation (gray shaded boxes).

**Supplementary Figure S5:**
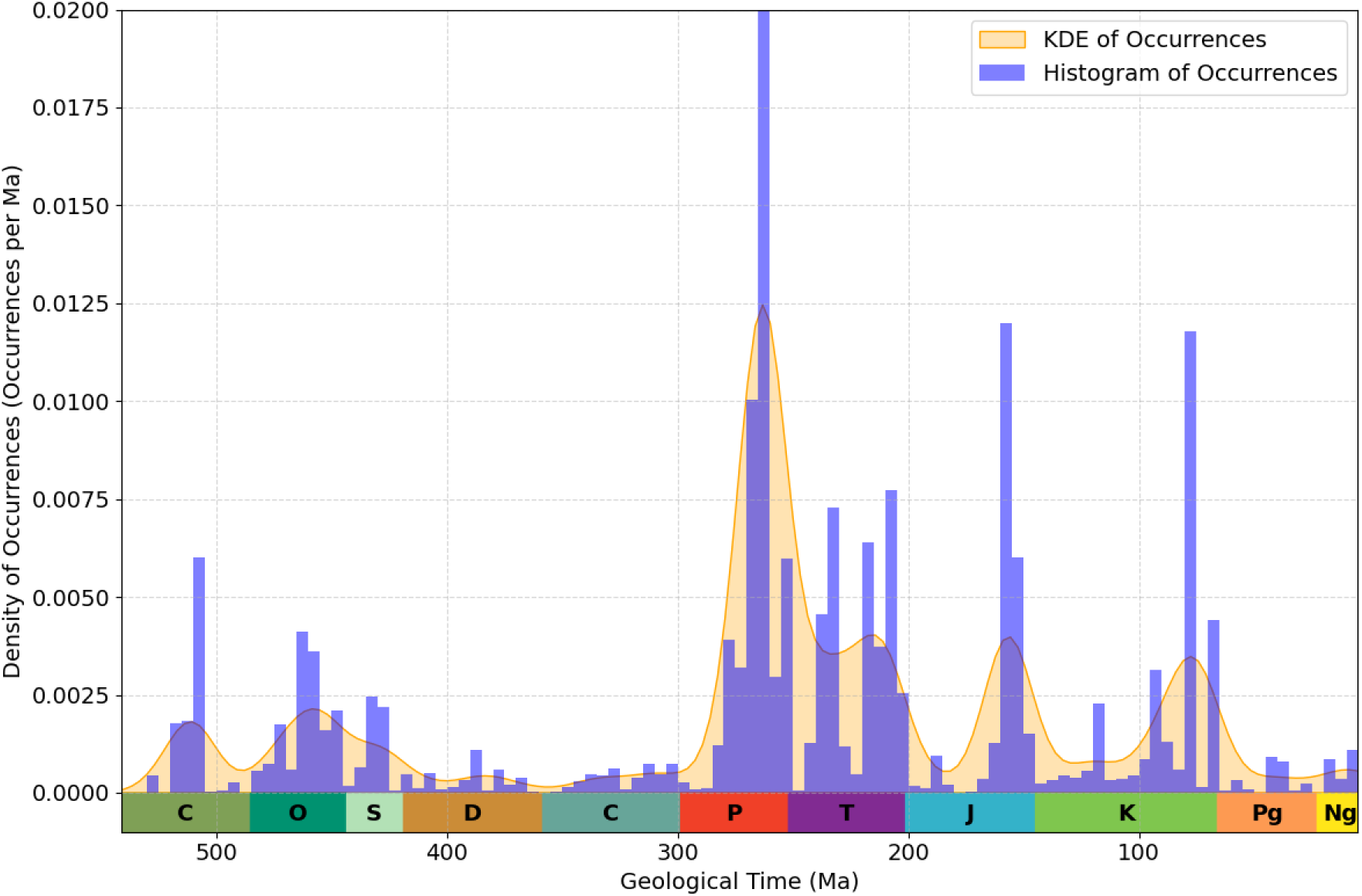
Density of fossil occurrences for Demospongiae in the Phanerozoic (541–5 Ma), based on the Paleobiology Dataset entry in PyRate. The plot integrates a Kernel Density Estimate (KDE) curve (orange) and a histogram (blue) showing the density of fossil occurrences using a 5-million-year time bin. The y-axis indicates normalized occurrence density.

**Supplementary Figure S6:**
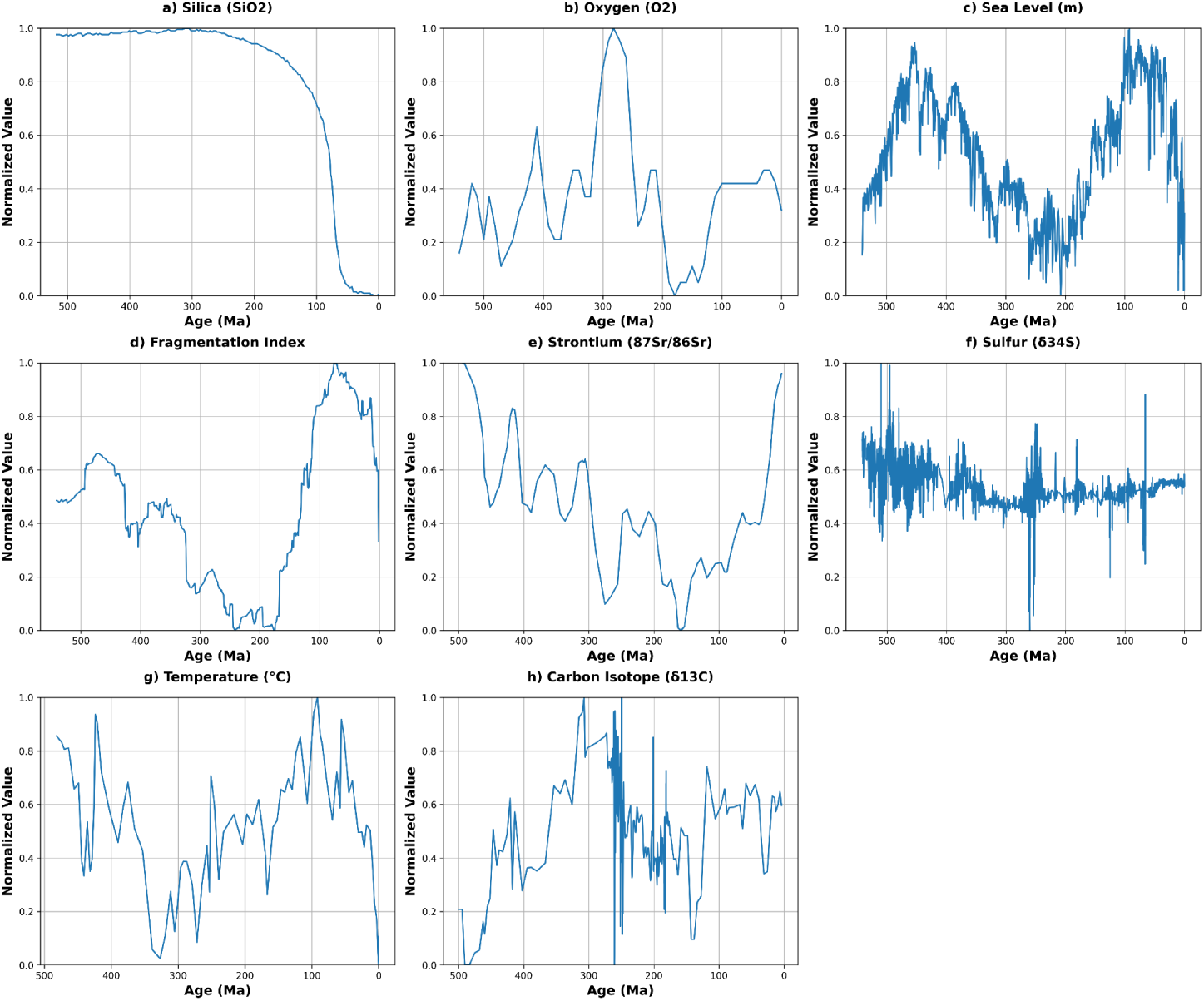
Normalized trends of key geological and environmental indicators over the Phanerozoic used to estimate correlation parameters on origination and extinction rates with abiotic factors (see. Fig. 3). Each subplot represents the following parameters: a) Silica (SiO2) concentration (Conley et al. 2017), b) Oxygen (O2) levels (Berner 2009; Lenton et al. 2018), c) Sea Level changes (Haq and Schutter 2008; Haq 2018), d) Fragmentation Index reflecting global continental fragmentation (Zaffos et al. 2017), e) Strontium (87Sr/86Sr) isotope composition (McArthur et al. 2020), f) Sulfur (δ^34^S) isotope composition (Present et al. 2020), g) Temperature (°C) variations estimated from marine δ18O (Scotese et al. 2021), and h) Carbon Isotope (δ13C) composition (Cramer and Jarvis 2020). Data are normalized using the formula (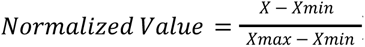) for each variable where *X* is the observed value, and *Xmin* and *Xmax* are the minimum and maximum values, respectively, for each dataset to fit within a range of 0 to 1 for comparison across different scales. The x-axis represents age in millions of years ago (Ma).

**Supplementary Fig. S7.**
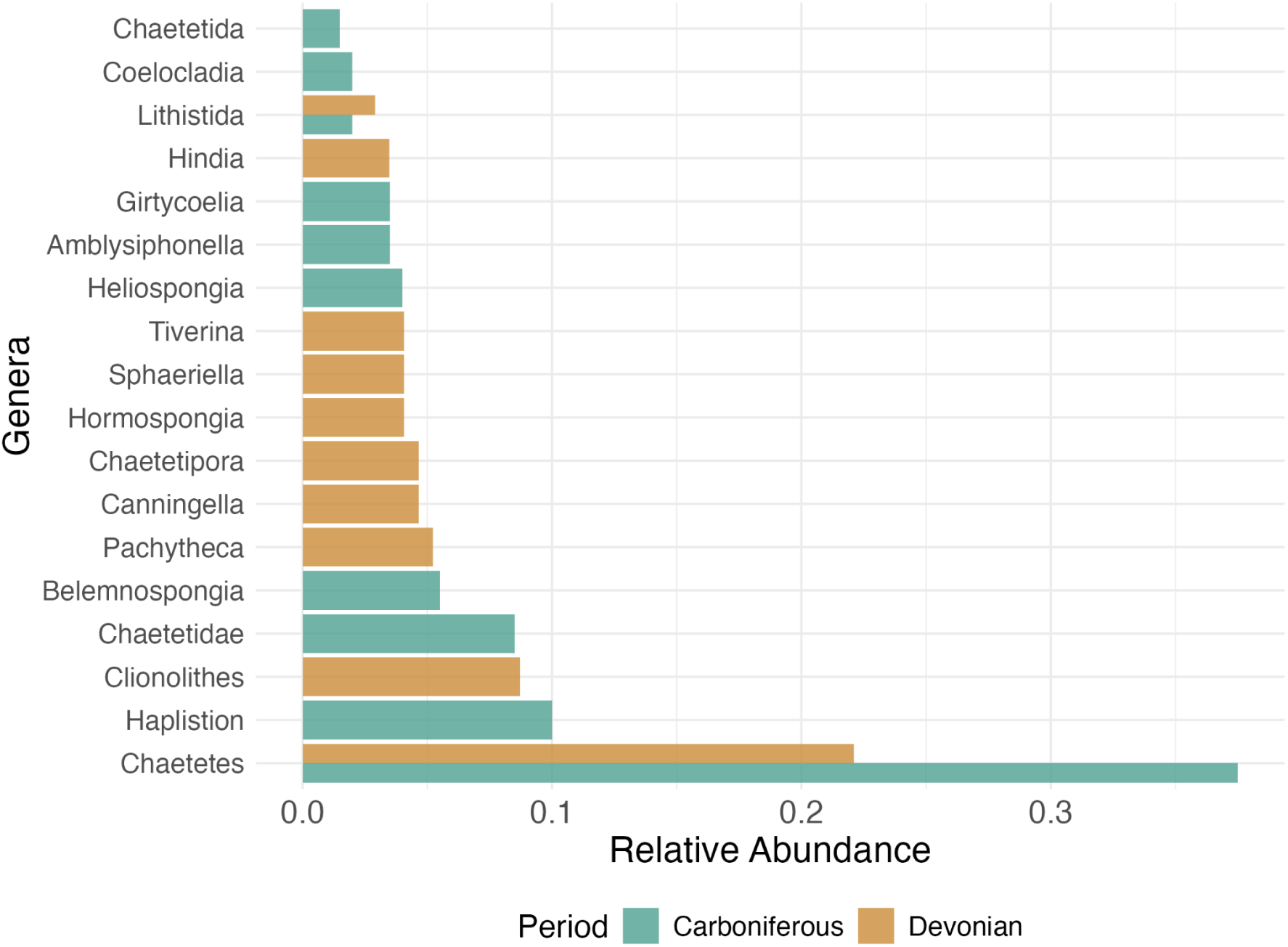
Relative abundance of top 10 dominant genera/taxa in the Dvonian and Carboniferous Period. The plot shows the proportion of each genus/taxa within these two periods.

